# One year prevalence of psychotic disorders among first treatment contact patients at Butabika National Psychiatric Referral Hospital in Uganda

**DOI:** 10.1101/669606

**Authors:** Emmanuel Kiiza Mwesiga, Noeline Nakasujja, Juliet Nakku, Annet Nanyonga, Joy Louise Gumikiriza, Paul Bangirana, Dickens Akena, Seggane Musisi

## Abstract

**Introduction:** Hospital based studies for psychotic disorders are scarce in low and middle income countries. This may impact on development of intervention programs.

**Objective:** We aimed to determine the burden of psychotic disorders among first treatment contact patients at the national psychiatric referral hospital in Uganda.

**Methods:** A retrospective patient chart-file review was carried out in March 2019 for all patients presenting to the hospital for the first time in the previous year. Patients were categorised into those with and without psychotic disorders. We collected sociodemographic data on age, gender, occupation, level of education, ethnicity, religion and home district. We determined the one year prevalence of psychotic disorders among first treatment contact patients. Using logistic regression models, we also determined the association between psychotic disorders and various exposure variables among first treatment contact patients.

**Results:** In 2018, 63% (95% CI: 60.2 – 65.1) of all first time contact patients had a psychosis related diagnosis. Among the patients with psychotic disorders, the median age was 29 years (IQR 24 – 36). Most of the patients were male (62.8%) and unemployed (63.1%). After adjusting for patients’ residence, psychotic disorders were found to be more prevalent among the female gender [OR 1.58 (CI1.46-1.72)] and those of Pentecostal faith [OR 1.25 (CI 1.10-1.42)].

**Conclusion:** Among first treatment contact patients in Uganda, there is a large burden of psychotic disorders. The burden was more prevalent among females as well as people of Pentecostal faith who seemed to use their church for faith-based healing. Incidence studies are warranted to determine if this phenomenon is replicated at illness onset.

## INTRODUCTION

Psychotic disorders that include schizophrenia spectrum disorders as well as bipolar affective disorders are the leading contributors to disease burden globally (1–3). Schizophrenia was assigned the highest disability coefficient in global burden of disease (GBD) study (4, 5). Psychotic disorders run a chronic course in the life of an individual. They usually present in early adolescence with a first episode of psychosis; and then continue with some form of disability thorough out the life of the individual (6). Patients with psychotic disorders are more likely to have worse social functioning, poor quality of life and die earlier than their peers (7–12). Correct management at initial presentation of psychotic disorders has been associated with lower relapse rates, greater functional recovery and improved quality of life (13, 14). Worldwide the prevalence for psychotic disorders has remained relatively stable between 1-3% even in low and middle income countries (LMIC) like Uganda (3). Hospital based prevalence rates for psychotic disorders especially among first time attended in LMIC are however scarce. The current literature in the Ugandan setting has mainly dwelt on people with HIV/AIDS among first time mental treatment contacts (15).

There is limited literature on the burden of psychotic disorders at initial mental treatment contact in LMICs (16). It is unclear if the burden of psychotic disorders is greater than that for other disorders like anxiety, mood or substance use disorders. Such information is crucial in human resource allocation and the development of specialised services in tertiary care. The sociodemographic profile of patients presenting to tertiary care in the Ugandan setting is not well described. For example, literature has shown higher incident rates for psychotic disorders among males than females (17–21). Whether this is replicated at presentation for care in our setting is unknown. Also, the clinical profiles of the various psychotic disorders are unknown. This is especially important as management differs between the different psychosis spectrum disorders (22). The majority of patients with psychotic disorders prefer alternative and complimentary therapies over western medicine (23–29). It is unclear if this preference translates to lower rates and/or different clinical profiles for psychotic disorders among patients presenting to mental health services for the first time. Such differences are important in directing policy and developing interventions to improve care for patients with psychotic disorders.

Describing the burden and risk factors for psychotic disorders at initial treatment contact is a crucial step in developing interventions to improve the outcomes for patients with psychotic disorders. In Uganda there is a precedent for this approach where extensive literature on the burden of HIV/AIDS in the psychiatric setting was instrumental in development of interventions for patients with severe mental illness suffering with AIDS (30–34). The current study therefore aims to determine the burden of psychotic disorders among initial treatment contact patients at the national psychiatric hospital in Uganda.

## METHODS

The study took place at Butabika National Psychiatric Referral and Teaching Hospital, a 600 bed capacity mental hospital located approximately twelve kilometres from Kampala (35). The hospital is located in the heart of the Greater Kampala Metropolitan Area (GKMA) where 10% of Uganda’s population reside and responsible for a third of the country’s gross domestic product (GDP) (36). Butabika National Psychiatric Referral and Teaching Hospital determines the policy agenda for mental health in the country together with the Ministry Of Health and is responsible for various levels of mental health training (37). It also plays a supervisory role over all mental health provision services in the country that include 12 regional referral hospitals and 96 district hospitals. Functioning below the district hospitals are three different levels of health centres (HC) namely HC4, HC3 and HC2. Mental health provision starts at HC3 level with subsequent referrals to higher centres. Currently, the hospital has specialised services for substance use disorders at the Alcohol and drug unit, a forensic ward, a specialised child and adolescent mental health unit as well as specialised occupational therapy and psychotherapy units. In terms of human resource allocation, the national psychiatric and teaching hospital is run by 72 clinicians (psychiatrists’ clinical psychologists and psychiatric clinical officers); 157 nurses, 4 social workers and 59 mental attendants. Given that it is a national referral hospital it also provides non psychiatric care like HIV/AIDS care, minor surgeries and dental services. Like in many similar facilities in LMICs there are a number of challenges in provision of services primarily due to limited budgetary allocation (37, 38).

We used a retrospective case analysis of chart records to determine the burden, profile and associated factors for psychotic disorders among first treatment contact patients. Approval for the study was obtained from the Uganda National Council for Science and Technology (UNCST) and the School of Medicine Research and Ethics Committee (SOMREC) of Makerere University. We also received institutional approval from the hospital to carry out the study. As this was a retrospective chart review of file records, we did not receive patient consent. All patients presenting to the hospital for the first time who had a psychiatric diagnosis on file between January 1^st^ and December 31^st^, 2018 made our study population. We excluded patients presenting for the first time for non-psychiatric services like dental services, routine HIV care or minor surgeries like circumcision.

On a routine clinic day, the hospital records team opens a file for all patients presenting to the hospital for the first time. The patient sociodemographic variables including age, gender, ethnicity, religion, occupation and home district are recorded in the file before the patient is sent to see a clinician. The clinician then makes a diagnosis, and a decision of whether to treat the patient as an out-patient or send them to admission in one of the units described above. Once the patient has received care, the health care workers return the patient file to the records office for safe storage. Some patients receive care as in-patients, and others are treated as out-patients and return to their homes the same day.

We used standardized questionnaires to extract sociodemographic and diagnosis data from the chart files of all patients presenting to the hospital for the first time from January to December 2018. Diagnoses of schizophrenia spectrum and related psychoses, bipolar affective disorder and mood disorders with psychotic disorders were classified as psychotic disorders. All other diagnoses among patients presenting for the first time including but not limited to temporal lobe epilepsy, anxiety disorders, substance use disorders and depressive disorders were classified as non-psychotic disorders. We considered sociodemographic characteristics as the exposure variables and the diagnostic categories as the outcome variables. Abstracted data from the files was entered into Epidata 3.1 by a database manager and exported to Stata version 13 for analysis. Data analysis was conducted in March 2019.

Proportions of patients by different diagnostic categories were calculated to determine the one year prevalence of psychotic disorders. Using bivariate analysis we compared the proportions of participants with psychotic disorders to non-psychotic disorders along various exposures. No variables exhibited any collinearity and the dataset had no outliers. We used a modified Poisson regression model to establish factors associated with psychotic disorders given that it has robust standard errors and therefore gives more accurate confidence intervals. Variables with a level of significance less than 0.2 were included in the multivariate analysis. However, region of origin was assessed for any possible confounding effects as ethnicity has been shown to have a genetic biological risk factor for psychotic disorders At multi-variate analysis a level of significance of less than 0.05 was used to test for significance between different exposures and FEP.

## RESULTS

Between January 1^st^, 2018 and December 31^st^, 2018; 1685 patients accessed services from Butabika for the first time. A total of 201 (11.93%) patients lacked a diagnosis in their records and were excluded from the final analysis. The total number of records reviewed for this study was 1484. On average there were 5 new patients each day accessing the hospital for the first time during the year 2018. Figure 1 shows the proportions of patients seen by month and gender. Other baseline characteristics of all new participants are highlighted in table 1. Among all new patients, the commonest diagnosis was a non-affective psychosis accounting for 32.01% of the total sample closely followed by substance use disorder at 30.39%. Anxiety disorders were the least common final diagnosis at 0.47%. The frequencies of different diagnoses among the total sample are highlighted in Figure 2.

**Table 1:** Background characteristics of all patients who reported for the first time in 2018.

| Variable | Number (N) | Percentage (%) |
| --- | --- | --- |
| ≤ 29 | 757 | 52.9 |
| > 29 | 674 | 47.1 |
| **Gender** |  |  |
| Male | 930 | 62.8 |
| Female | 549 | 37.1 |
| **Religion** |  |  |
| Protestant | 404 | 32.3 |
| Catholic | 407 | 32.5 |
| Moslem | 242 | 19.3 |
| Seventh day Adventist | 29 | 2.3 |
| Pentecostal/Born again | 123 | 9.8 |
| Other religions | 46 | 3.7 |
| **Occupation** |  |  |
| Student | 89 | 6.4 |
| Formal | 108 | 7.8 |
| Non-formal | 270 | 19.4 |
| Unemployed | 922 | 66.4 |
| **Region** |  |  |
| Central | 1,093 | 79.9 |
| Eastern | 102 | 7.5 |
| Northern | 30 | 2.2 |
| Western | 143 | 10.5 |

**Figure 1:**
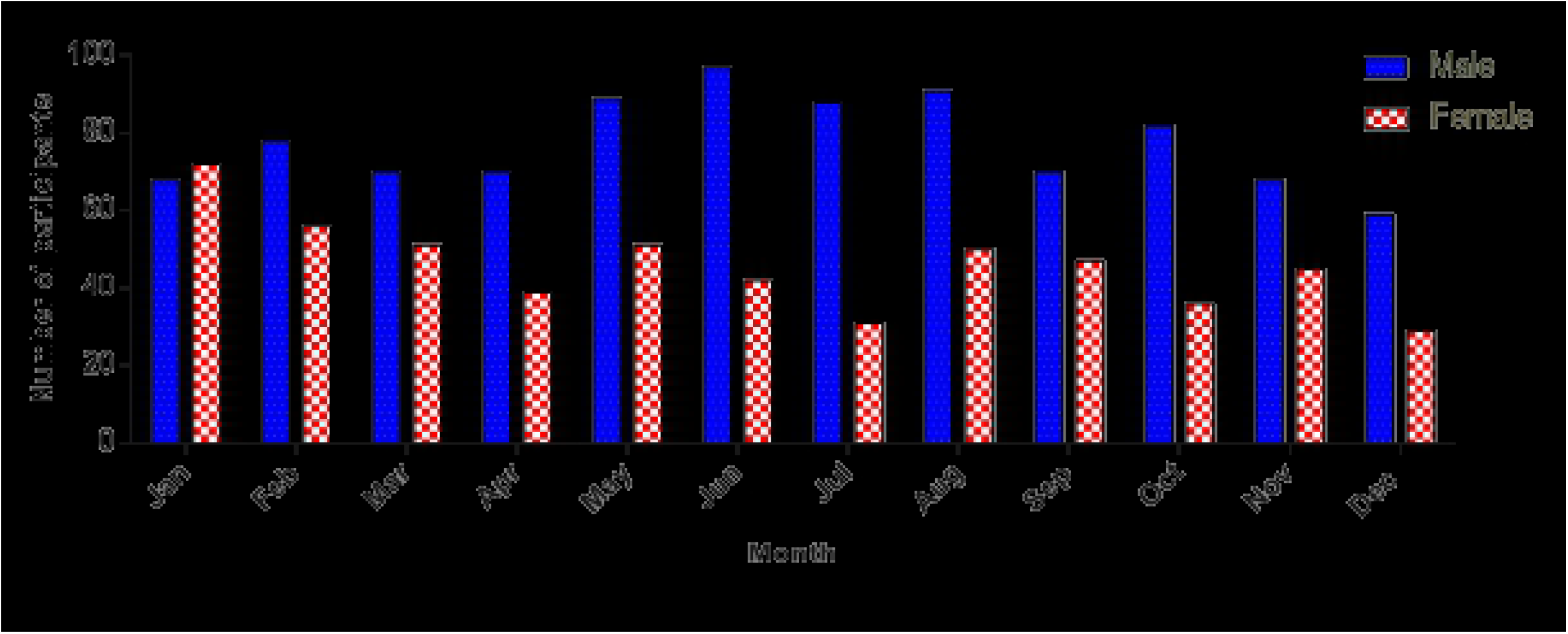
Bar Graph of number of participants by month of the year and gender

**Figure 2:**
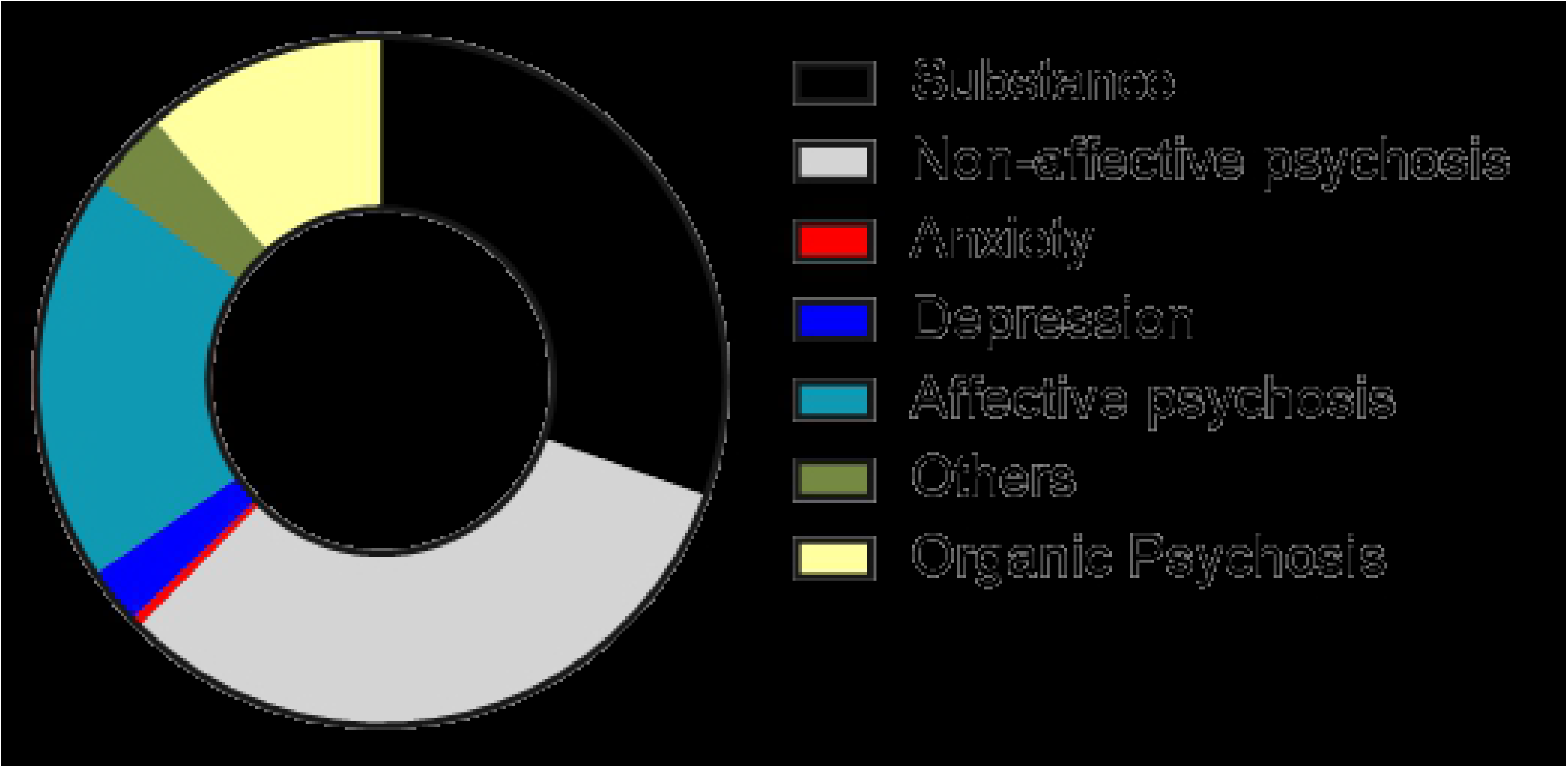
A pie chart showing the different diagnostic categories for the whole sample.

### Burden of psychotic disorders

Approximately two-thirds [62.7% (95% CI: 60.2 – 65.1)] of all patients had a psychotic disorder. Among the patients classified as having psychotic disorders, 51.08% were classified as having schizophrenia spectrum disorders, 30.75% as bipolar affective disorders and 18.17% as an organic psychosis. The median age for patients with psychotic disorders was 29 years (IQR 24 – 36) with almost twice as many males as females. Most participants (76.03%) were between the 30 to 39 age range with only 4.54% of patients below the age of 18 years. Other baseline characteristics of the patients with psychotic disorders are shown in table 2.

**Table 2:**
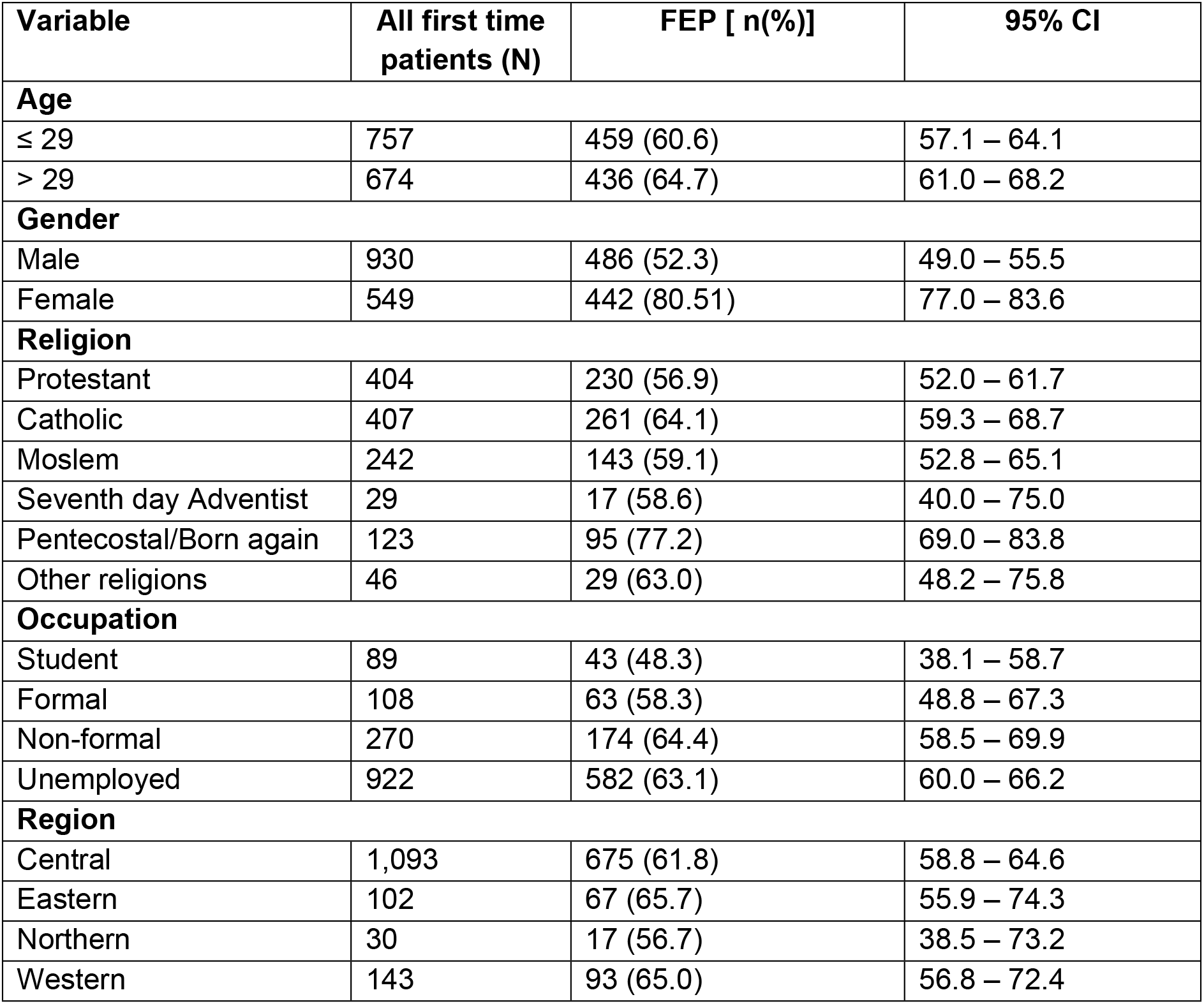
Background characteristics of the sample of participants classified as having psychosis.

At bi-variate analysis, psychotic disorders were found to be more prevalent among the female gender [Prevalence ratio (PR) 1.54 (confidence interval 1.43-1.66)] as well as patients who reported to subscribe to the Catholic [PR 1.13 (CI 1.01-1.26)] or Pentecostal faiths [PR 1.36 (CI 1.19-1.54)]. Psychotic disorders were also more prevalent among patients of non-formal employment, the unemployed as well as those presenting in the month of November. Other associations are highlighted in Table 3.

**Table 3:** Bivariate analysis of the association between patients with a psychosis diagnosis and different sociodemographic variables.

| Variable | Total (N) | FEP<br>Prevalence<br>n(%) | Prevalence<br>ratio | 95% CI | P-value |
| --- | --- | --- | --- | --- | --- |
| Age |  |  |  |  |  |
| ≤ 29 | 757 | 459 (60.6) | 1 | 0.98 – 1.16 | 0.113 |
| > 29 | 674 | 436 (64.7) | 1.07 |  |  |
| Gender |  |  |  |  |  |
| Male | 930 | 486 (52.3) | 1.00 | 1.43 – 1.66 | < 0.001 |
| Female | 549 | 442 (80.51) | 1.54 |  |  |
| Religion |  |  |  |  |  |
| Protestant | 404 | 230 (56.9) | 1.00 |  |  |
| Catholic | 407 | 261 (64.1) | 1.13 | 1.01 – 1.26 | 0.037 |
| Moslem | 242 | 143 (59.1) | 1.04 | 0.91 – 1.19 | 0.588 |
| Seventh day<br>Adventist | 29 | 17 (58.6) | 1.03 | 0.75 – 1.41 | 0.857 |
| Pentecostal/Born<br>again | 123 | 95 (77.2) | 1.36 | 1.19 – 1.54 | < 0.001 |
| Other religions | 46 | 29 (63.0) | 1.11 | 0.87 – 1.40 | 0.399 |
| Occupation |  |  |  |  |  |
| Student | 89 | 43 (48.3) | 1.00 |  |  |
| Formal | 108 | 63 (58.3) | 1.21 | 0.92 – 1.58 | 0.168 |
| Non-formal | 270 | 174 (64.4) | 1.33 | 1.06 – 1.68 | 0.015 |
| Unemployed | 922 | 582 (63.1) | 1.31 | 1.05 – 1.63 | 0.018 |
| Region |  |  |  |  |  |
| Central | 1,093 | 675 (61.8) | 1.00 |  |  |
| Eastern | 102 | 67 (65.7) | 1.06 | 0.92 – 1.23 | 0.414 |
| Northern | 30 | 17 (56.7) | 0.92 | 0.67 – 1.26 | 0.594 |
| Western | 143 | 93 (65.0) | 1.05 | 0.93 – 1.20 | 0.4321 |

In the final multi-variate model, gender [Prevalence ratio (PR) 1.58 (confidence interval 1.46-1.72)], and Pentecostal faith [PR1.25 (CI1.10-1.42)] remained significant after controlling for the region of the country the patient was from. Other associations are highlighted in table 4.

**Table 4:** Multivariate analysis of the association between FEP and selected exposures.

| Variable | Prevalence ratio | 95% CI | P-value |
| --- | --- | --- | --- |
| Age |  |  |  |
| ≤ 29 | 1.00 | 0.92 – 1.09 | 0.971 |
| > 29 | 0.99 |  |  |
| Gender |  |  |  |
| Male | 1.00 | 1.46 – 1.72 | < 0.001 |
| Female | 1.58 |  |  |
| Religion |  |  |  |
| Protestant | 1.00 |  |  |
| Catholic | 1.11 | 1.00 – 1.24 | 0.050 |
| Moslem | 1.04 | 0.91 – 1.18 | 0.603 |
| Seventh day Adventist | 1.00 | 0.74 – 1.36 | 0.857 |
| Pentecostal/Born again | 1.25 | 1.10 – 1.42 | 0.001 |
| Other religions | 1.14 | 0.87 – 1.48 | 0.340 |

## DISCUSSION

### Mental health service requirements for patients with psychotic disorders

Over two-thirds (67%) of all admissions presenting to the hospital for the first time in 2018 had a psychotic disorder. To our knowledge this is the first published study highlighting the large burden of psychotic disorders in the Ugandan setting among patients presenting for the first time at a mental facility. The burden for psychotic disorders was greater than that for mood disorders as well as substance use disorders. This suggests that there may be benefit in introducing specialised early intervention services for psychotic disorders at the hospital. Specialised services for psychotic disorders especially at the first episode of psychosis usually lead to better outcomes for patients (39–42). Currently the hospital has specialised services for substance use disorders, and it would be important to determine the benefit of similar services for psychotic disorders. Future work on necessary components for an early intervention psychosis clinic as well as cost benefit analyses of such a program are recommended (13, 21, 42, 43). It is also known and often observed that psychotic disorders tend to present with aggression and violence injuring staff and fellow patients (44, 45). Acute psychiatric units or psychiatric intensive care units have been shown to be especially effective in containing such potentially dangerous behaviour (44), hence calling for such care facilities as useful additions to mental hospitals as opposed to just locked seclusion rooms as is the practice at this facility (44, 45).

### Time of presentation and duration of untreated illness

The low numbers of patients presenting to the hospital younger than 18 years of age is worrying as it may point to delay in presentation for services. The course of psychotic disorders is characterised by a psychosis prodrome before onset of illness usually in the late teens or early adulthood (40, 46). That most of our patients present outside this age range may imply that either the onset of psychosis is late in this population or that there is a long duration of untreated psychosis (DUP). The latter theory is probably more likely since DUP has been reported to be longer in Sub-Saharan Africa compared to high income countries (47–49). This is important for future intervention programs given that DUP is a key predictor of outcomes for patients with psychotic disorders (14, 46, 50).

### Gender and initial presentation to care with psychotic disorders

Females were more likely to present to the hospital than males with a psychotic illness. The incidence of psychotic disorders is higher in males than females in previous literature (17–21). Greater prevalence among the female gender might be due to the difference in care seeking between males and females rather than greater incidence in the community. This, however, would need confirmation with longitudinal studies. It is also important to note that it is unlikely that a patient with psychosis brought themselves to the hospital. Further studies are therefore required to understand why there is preference for bringing females to the hospital than males.

### Culture and initial presentation to care with a psychotic disorder

Culture plays an important role in symptom presentation, care seeking and access to health services (51, 52). From this study, it is not possible to determine why there is greater prevalence for initial presentation at the hospital for psychotic disorders over non-psychotic disorders. Previous literature by Abbo et al (2009) highlighted that patients are more likely to use both African traditional therapies and biomedicine if the patient has a severe illness or poor global functioning (23). It is therefore possible that the patients coming to the hospital are the ones who were very ill and generally disruptive in the communities in which they lived. Unfortunately, this chart review could not answer this question but further highlights that patients may be coming late with long duration of untreated psychosis. Previous literature has highlighted the preference for alternative and complementary therapies for the initial management of psychotic disorders in this setting (23, 24, 26, 27).

Psychotic disorders were more prevalent among people of the Pentecostal faith. It is important to clarify that this finding does not mean that people of this faith are more at risk for psychotic disorders. Rather the findings suggest that people of Pentecostal faith with psychotic disorders were more likely than other faiths to seek care from the national referral and psychiatric hospital. Another plausible explanation might be due to explanatory models for mental illness in our setting characterised by beliefs in supernatural causations of psychotic disorders (53). This may make patients resort to this faith because of its supposed ability to heal mental disorders through prayer hence leading to more psychotic cases there eventually presenting to the hospital (54, 55).

Ethnicity has a strong association to genetic risk which is a key biological risk factor for psychotic disorders (56, 57). Psychotic disorders were not found to be more prevalent in any particular ethnic grouping or region of origin. Uganda is one of the most ethnically diverse societies in the world (58) and this sample had more than 30 different tribes. It would therefore require larger sample sizes to determine an association between a specific ethnicity and onset of psychotic disorders. Currently a large genetic study is underway in Uganda to try and determine the genetic risk for psychotic disorders (59).

### Limitations of the study

A major limitation of the study was its retrospective study design which could cause information bias. The information however collected was primarily on sociodemographic characteristics which are not usually prone to bias. Also, failure to confirm the diagnoses with a standardized tool could lead to misclassification bias. However, Butabika is a national referral hospital with expertise in mental health care service provision and the diagnoses were made by qualified psychiatrists; so we were fairly confident in the diagnoses made.

## CONCLUSION

There seems to be a large burden of psychotic disorders (67%) among patients presenting to the national psychiatric hospital in Uganda for the first time. Many of the participants were female calling for further studies to understand this phenomenon in our setting. More studies are also needed to define the duration of untreated psychosis in this population given that most of the first time patients were older than the normal onset for psychotic disorders. Finally, there may be benefits in introducing specialised intervention services for psychotic disorders at the national referral hospital in the form of specialised early intervention services as well as “safe wards models” as acute psychiatric units or psychiatric intensive care units at such large mental health facilities

## ACKNOWLEDGEMENT

We acknowledge the patients who presented to the hospital for the first time. Dr. Linnet Ongeri of Kenya Medical Research Institute gave invaluable guidance on the manuscript for which we are grateful.

## Funding

The work was supported by Grant Number D43TW010132 supported by Office Of The Director, National Institutes Of Health (OD), National Institute Of Dental & Craniofacial Research (NIDCR), National Institute Of Neurological Disorders And Stroke (NINDS), National Heart, Lung, And Blood Institute (NHLBI), Fogarty International Centre (FIC), National Institute On Minority Health And Health Disparities (NIMHD). Its contents are solely the responsibility of the authors and do not necessarily represent the official views of the supporting offices.

## Competing Interests

The authors declare no competing interests.

## Author contributions

EKM, NN and SM conceptualised the research idea. EKM, AN, JN and JLG supervised the data extraction exercise. PB and DA advised on the analysis of the results. All authors were involved in writing the manuscript and approved the final manuscript for submission.

## Data Availability

The data underlying the results presented in the study are available from the corresponding author on request.

## REFERENCES

1. Hyman S, Parikh R, Collins PY, Patel V. Adult Mental Disorders.

2. Ruggeri M, Leese M, Thornicroft G, Bisoffi G, Tansella M. Definition and prevalence of severe and persistent mental illness. The British Journal of Psychiatry. 2000;177(2):149–55.

3. Association AP. DSM 5: American Psychiatric Association; 2013.

4. Rössler W, Salize HJ, van Os J, Riecher-Rössler A. Size of burden of schizophrenia and psychotic disorders. European Neuropsychopharmacology. 2005;15(4):399–409.

5. Salomon JA, Vos T, Hogan DR, Gagnon M, Naghavi M, Mokdad A, et al. Common values in assessing health outcomes from disease and injury: disability weights measurement study for the Global Burden of Disease Study 2010. Lancet. 2012;380(9859):2129–43.

6. Urs Heilbronner, Myrto Samara, Stefan Leucht, Peter Falkai, Thomas G. Schulze. The Longitudinal Course of Schizophrenia Across the Lifespan: Clinical, Cognitive, and Neurobiological Aspects. Harvard Review of Psychiatry. 2016;24(2):118–28. doi:10.1097/HRP.0000000000000092.

7. Broussard B, Kelley ME, Wan CR, Cristofaro SL, Crisafio A, Haggard PJ, et al. Demographic, socio-environmental, and substance-related predictors of duration of untreated psychosis (DUP). Schizophrenia Research. 2013;148(1):93–8.

8. Emsley R, Chiliza B, Schoeman R. Predictors of long-term outcome in schizophrenia. Curr Opin Psychiatry. 2008;21(2):173–7.

9. Erin E Michalak, Lakshmi N Yatham, Raymond W Lam. Quality of life in bipolar disorder: A review of the literature. Health and quality of life outcomes. 2005;3(72):doi:10.1186/477-7525-3-72.

10. Abebaw Fekadu, Girmay Medhin, Derege Kebede, Atalay Alem Excess mortality in severe mental illness: 10-year population-based cohort study in rural Ethiopia. The British Journal of Psychiatry. 2015;206:289–96. doi: 10.1192/bjp.bp.114.149112.

11. Gregory E. Simon, Christine Stewart, Bobbi Jo Yarborough, Frances Lynch, Karen J. Coleman, Arne Beck, et al. Mortality Rates After the First Diagnosis of Psychotic Disorder in Adolescents and Young Adults. JAMA Psychiatry. 2018;doi:10.1001/jamapsychiatry.2017.4437.

12. Solomon Teferra, Teshome Shibre, Abebaw Fekadu, Girmay Medhin, Asfaw Wakwoya, Atalay Alem, et al. Five-year mortality in a cohort of people with schizophrenia in Ethiopia. BMC Psychiatry,. 2011;11::165 doi.org/10.1186/471-244X-11-165.

13. Marshall M, Lockwood A, Lewis S, Fiander M. Essential elements of an early intervention service for psychosis: the opinions of expert clinicians. BMC Psychiatry. 2004;4:17.

14. Marshall M, Rathbone J. Early intervention for psychosis. (1469-493X (Electronic)).

15. Maling S, Todd J, Van der Paal L, Grosskurth H, Kinyanda E. HIV-1 seroprevalence and risk factors for HIV infection among first-time psychiatric admissions in Uganda. AIDS care. 2011;23(2):171–8.

16. Sharifi V, Amin-Esmaeili M, Hajebi A, Motevalian A, Radgoodarzi R, Hefazi M, et al. Twelve-month prevalence and correlates of psychiatric disorders in Iran: the Iranian Mental Health Survey, 2011. Archives of Iranian medicine. 2015;18(2):76–84.

17. Anderson KK, Fuhrer R, Abrahamowicz M, Malla AK. The incidence of first-episode schizophrenia-spectrum psychosis in adolescents and young adults in montreal: an estimate from an administrative claims database. Can J Psychiatry. 2012;57(10):626–33.

18. Anderson KK, Norman R, MacDougall AG, Edwards J, Palaniyappan L, Lau C, et al. Estimating the incidence of first-episode psychosis using population-based health administrative data to inform early psychosis intervention services. Psychol Med. 2018:1–9.

19. Elliot M Goldner, Lorena Hsu, Paul Waraich, Julian M Somers. Prevalence and Incidence Studies of Schizophrenic Disorders: A Systematic Review of the Literature. Can J Psychiatry. 2002;47:833–43.

20. Hannah E. Jongsma, Charlotte Gayer-Anderson, Antonio Lasalvia, Diego Quattrone, Alice Mulè, Andrei Szöke, et al. Treated Incidence of Psychotic Disorders in the Multinational EU-GEI Study. JAMA Psychiatry. 2018;JAMA Psychiatry(75):1.

21. Perez J, Russo DA, Stochl J, Shelley GF, Crane CM, Painter M, et al. Understanding causes of and developing effective interventions for schizophrenia and other psychoses.

22. Kane JM, Leucht S, Carpenter D, Docherty JP. The expert consensus guideline series. Optimizing pharmacologic treatment of psychotic disorders. Introduction: methods, commentary, and summary. J Clin Psychiatry. 2003;64 Suppl 12:5–19.

23. Abbo C, Ekblad S, Waako P, Okello E, Musisi S. The prevalence and severity of mental illnesses handled by traditional healers in two districts in Uganda. Afr Health Sci. 2009;1(9):S16–22.

24. Abbo C, Odokonyero R, Ovuga E. A narrative analysis of the link between modern medicine and traditional medicine in Africa: a case of mental health in Uganda. Brain research bulletin. 2018.

25. Sorsdahl K, Stein DJ, Grimsrud A, Seedat S, Flisher AJ, Williams DR, et al. Traditional healers in the treatment of common mental disorders in South Africa. The Journal of nervous and mental disease. 2009;197(6):434–41.

26. Abbo C, Ekblad S, Waako P, Okello E, Muhwezi W, Musisi S. Psychological distress and associated factors among the attendees of traditional healing practices in Jinja and Iganga districts, Eastern Uganda: a cross-sectional study. Int J Ment Health Syst. 2008;2(1):1752–4458.

27. Abbo C, Okello ES, Musisi S, Waako P, Ekblad S. Naturalistic outcome of treatment of psychosis by traditional healers in Jinja and Iganga districts, Eastern Uganda - a 3- and 6 months follow up. Int J Ment Health Syst. 2012;6(1):1752–4458.

28. Schmid GB, Brunisholz K. [Evaluation of use of complementary and alternative medicine by schizophrenic patients]. Forschende Komplementarmedizin (2006). 2007;14(3):167–72.

29. Abbo C, Ekblad S, Waako P, Okello E, Musisi S. The prevalence and severity of mental illnesses handled by traditional healers in two districts in Uganda. Afr Health Sci. 2009;9 Suppl 1:S16–22.

30. Nakasujja N, Allebeck P, Agren H, Musisi S, Katabira E. Cognitive dysfunction among HIV positive and HIV negative patients with psychosis in Uganda. PLoS ONE. 2012;7(9).

31. Nakimuli-Mpungu E, Musisi S, Wamala K, Okello J, Ndyanabangi S, Mojtabai R, et al. The Effect of Group Support Psychotherapy Delivered by Trained Lay Health Workers for Depression Treatment Among People with HIV in Uganda: Protocol of a Pragmatic, Cluster Randomized Trial. JMIR research protocols. 2017;6(12):e250.

32. Akena D, Joska J, Stein DJ. Sensitivity and specificity of the Akena Visual Depression Inventory (AViDI-18) in Kampala (Uganda) and Cape Town (South Africa). Br J Psychiatry. 2018;212(5):301–7.

33. Kinyanda E, Salisbury TT, Levin J, Nakasujja N, Mpango RS, Abbo C, et al. Rates, types and co-occurrence of emotional and behavioural disorders among perinatally HIV-infected youth in Uganda: the CHAKA study. Soc Psychiatry Psychiatr Epidemiol. 2019;54(4):415–25.

34. Sacktor NC, Wong M, Nakasujja N, Skolasky RL, Selnes OA, Musisi S, et al. The International HIV Dementia Scale: a new rapid screening test for HIV dementia. AIDS (London, England). 2005;19(13):1367–74.

35. Kigozi F, Ssebunnya J, Kizza D, Cooper S, Ndyanabangi S. An overview of Uganda’s mental health care system: results from an assessment using the world health organization’s assessment instrument for mental health systems (WHO-AIMS). Int J Ment Health Syst. 2010;4(1):1.

36. Publication WB. GreatKampalaMetropolitanAreaQuickFacts.

37. Petersen I, Marais D, Abdulmalik J, Ahuja S, Alem A, Chisholm D, et al. Strengthening mental health system governance in six low- and middle-income countries in Africa and South Asia: challenges, needs and potential strategies. Health policy and planning. 2017;32(5):699–709.

38. Ssebunnya J, Kangere S, Mugisha J, Docrat S, Chisholm D, Lund C, et al. Potential strategies for sustainably financing mental health care in Uganda. Int J Ment Health Syst. 2018;12:74.

39. Coentre R, Levy P, Figueira ML. [Early intervention in psychosis: first-episode psychosis and critical period]. Acta medica portuguesa. 2011;24(1):117–26.

40. Moe AM, Rubinstein EB, Gallagher CJ, Weiss DM, Stewart A, Breitborde NJ. Improving access to specialized care for first-episode psychosis: an ecological model. Risk management and healthcare policy. 2018;11:127–38.

41. Rangaswamy T, Mangala R, Mohan G, Joseph J, John S. Early intervention for first-episode psychosis in India. East Asian archives of psychiatry: official journal of the Hong Kong College of Psychiatrists = Dong Ya jing shen ke xue zhi: Xianggang jing shen ke yi xue yuan qi kan. 2012;22(3):94–9.

42. Stafford MR, Jackson H, Mayo-Wilson E, Morrison AP, Kendall T. Early interventions to prevent psychosis: systematic review and meta-analysis. The BMJ. 2013;346:f185.

43. White DA, Luther L, Bonfils KA, Salyers MP. Essential components of early intervention programs for psychosis: Available intervention services in the United States. Schizophr Res. 2015;168(1-2):79–83.

44. Musisi SM, Wasylenki DA, Rapp MS. A psychiatric intensive care unit in a psychiatric hospital. Can J Psychiatry. 1989;34(3):200–4.

45. Baumgardt J, Jäckel D, Helber-Böhlen H, Stiehm N, Morgenstern K, Voigt A, et al. Preventing and Reducing Coercive Measures—An Evaluation of the Implementation of the Safewards Model in Two Locked Wards in Germany. Frontiers in Psychiatry. 2019;10(340).

46. Murru A, Carpiniello B. Duration of untreated illness as a key to early intervention in schizophrenia: A review. Neuroscience Letters. 2018;669:59–67.

47. Davis GP, Tomita A, Baumgartner JN, Mtshemla S, Nene S, King H, et al. Substance use and duration of untreated psychosis in KwaZulu-Natal, South Africa. The South African journal of psychiatry: SAJP: the journal of the Society of Psychiatrists of South Africa. 2016;22(1):a852.

48. Tomita A, Burns JK, King H, Baumgartner JN, Davis GP, Mtshemla S, et al. Duration of untreated psychosis and the pathway to care in KwaZulu-Natal, South Africa. J Nerv Ment Dis. 2015;203(3):222–5.

49. Chiliza B, Asmal L, Emsley R. Early intervention in schizophrenia in developing countries: focus on duration of untreated psychosis and remission as a treatment goal. Int Rev Psychiatry. 2012;24(5):483–8.

50. N. M. Menzes, T. Arenovich, R. B. Zipursky. A systematic review of longitudinal outcome studies of first-episode psychosis. Psychological Medicine. 2006;36:1349–62 doi:10.017/S0033291706007951.

51. Maraj A, Anderson KK, Flora N, Ferrari M, Archie S, McKenzie KJ. Symptom profiles and explanatory models of first-episode psychosis in African-, Caribbean- and European-origin groups in Ontario. Early Interv Psychiatry. 2017;11(2):165–70.

52. Singh SP, Brown L, Winsper C, Gajwani R, Islam Z, Jasani R, et al. Ethnicity and pathways to care during first episode psychosis: the role of cultural illness attributions. BMC Psychiatry. 2015;15:287.

53. Desai G, Chaturvedi SK. Idioms of Distress. Journal of neurosciences in rural practice. 2017;8(Suppl 1):S94–S7.

54. Kar N. Resort to faith-healing practices in the pathway to care for mental illness: a study on psychiatric inpatients in Orissa. Mental Health, Religion and Culture. 2008;11(7):720–40.

55. Peltzer K. Faith healing for mental and social disorders in the Northern Province (South Africa). Journal of Religion in Africa/Religion en Afrique. 1999;29(3):387.

56. Busby GB, Band G, Si Le Q, Jallow M, Bougama E, Mangano VD, et al. Admixture into and within sub-Saharan Africa. eLife. 2016;5.

57. Stevenson A, Akena D, Stroud RE, Atwoli L, Campbell MM, Chibnik LB, et al. Neuropsychiatric Genetics of African Populations-Psychosis (NeuroGAP-Psychosis): a case-control study protocol and GWAS in Ethiopia, Kenya, South Africa and Uganda. BMJ open. 2019;9(2):e025469.

58. Alesina AFaE, William and Devleeschauwer, Arnaud and Kurlat, Sergio and Wacziarg, Romain T.,. Fractionalization (June 2002). Harvard Institute Research Working Paper No. 1959.

59. Anne Stevenson, Dickens Akena, Rocky E Stroud, Lukoye Atwoli, Megan M Campbell, Lori B Chibnik, et al. Neuropsychiatric Genetics of African Populations-Psychosis (NeuroGAPPsychosis): a case-control study protocol and GWAS in Ethiopia, Kenya, South Africa and Uganda. BMJ open. 2019:e025469. doi:10.1136/bmjopen-2018-.

